# Postmating reproductive barriers contribute to the incipient sexual isolation of US and Caribbean *Drosophila melanogaster*

**DOI:** 10.1101/007765

**Authors:** Joyce Y. Kao, Seana Lymer, Sea H. Hwang, Albert Sung, Sergey V. Nuzhdin

**Author notes:** Contact Information: Joyce Kao, Sergey Nuzhdin. 1050 Childs Way, Los Angeles, CA, USA 90089-2910.

## Abstract

The nascent stages of speciation start with the emergence of sexual isolation. Understanding the influence of reproductive barriers in this evolutionary process is an ongoing effort. We present a study of *Drosophila melanogaster* populations from the southeast United States and Caribbean islands undergoing incipient sexual isolation. The existence of premating reproductive barriers have been previously established, but these types of barriers are not the only source shaping sexual isolation. To assess the influence of postmating barriers, we investigated putative postmating barriers of female remating and egg laying behavior, as well as hatchability of eggs laid and female longevity after mating. In the central region of our putative hybrid zone of American and Caribbean populations, we observed lower hatchability of eggs laid accompanied by increased resistance to harm after mating to less related males. These results illustrate that postmating reproductive barriers acting alongside premating barriers in a complex secondary contact zone. Furthermore, our findings suggest hybrid incompatibilities, likely due to the nature of genomic admixture of populations in the area, are influential even at the early phases of sexual isolation.

## Introduction

The onset of speciation is driven by reproductive barriers that reduce gene flow and result in reproductive isolation between populations. These barriers are classified by the temporal nature of their effect: prezygotic barriers occur before fertilization, while postzygotic barriers occur after fertilization (Coyne & Orr, 2004). The latter can be further divided into extrinsic and intrinsic categories, depending on whether the barrier interacts with external factors (e.g. environmental, individuals) or internal factors (e.g. genetic incompatibilities) (Seehausen *et al.*, 2014). Speciation involves multiple reproductive barriers of varying effect sizes (Coyne & Orr, 2004; Seehausen *et al.*, 2014), and identifying the interaction and strengths of reproductive barriers at play is vital to characterizing the process of speciation.

*Drosophila* is particularly well-suited to study reproductive barriers because this genus spans the whole speciation spectrum, from non-interbreeding species to hybridizing species (Bono & Markow, 2009) and populations (Yukilevich & True, 2008b). Empirical studies of sexual selection in *D. melanogaster* have investigated the evolution of prezygotic isolation - mate choice, male morphology, and courtship behavior (Hollocher, 1997; Yukilevich & True, 2008a). Postzygotic barrier mechanisms are also known to have an influence in *Drosophila*, but these studies have been limited to the hybridizing species *D. mojavensis/D. arizonae* (Bono & Markow, 2009) and *D. melanogaster/D. simulans* (Matute *et al.*, 2014).

Many natural forces influence the development of reproductive barriers; one example is sexual conflict, derived from the competing reproductive interests between males and females (Parker, 1979). Males may benefit from overriding the mating preferences evolved by females, and females consequently evolve resistance to these male ‘coercion’ tactics (Holland & Rice, 1998). Males are then selected for novel or more exaggerated traits - perpetuating an endless evolutionary chase between the sexes (Parker, 1979; Civetta & Singh, 1995; Rice, 1996; Chapman *et al.,* 2003; Arnqvist & Rowe, 2005; Arbuthnott *et al.,* 2014). This phenomenon of conflict in reproductive optima has been experimentally demonstrated to promote an antagonistic male-female coevolution that is the essence of sexual isolation which precedes speciation (Parker, 1979; Holland & Rice, 1998; Chapman *et al.*, 2003).

In *Drosophila melanogaster*, male sperm consists of accessory gland proteins that reduce female remating rates and increase egg laying (Chapman *et al.,* 2003; Wolfner, 1997). Reduced receptivity to remating will also decrease the female’s opportunity to mate with another male that could result in fitter progeny. Increased egg laying and the trauma from mating reduces female lifespan (Fowler & Partridge, 1989). As a result, females develop resistance to these harmful male traits, and males subsequently evolve new methods to discourage females from mating with other males (Arnqvist & Rowe, 2005). It has been suggested that females should be more resistant to males they have coevolved with (‘homotypic’) compared to males they have not coevolved with (‘heterotypic’). However, these effects vary across populations, and ecological context appears to be a factor (Arbuthnott *et al.*, 2014). This rapid, cyclical process termed sexually antagonistic coevolution has been demonstrated not only in *Drosophila* species (Knowles & Markow, 2001), but also in other organisms like water striders (Rowe & Arnqvist, 2002). Coevolution by sexual conflict is a strong force behind reproductive isolation, which may lead to speciation in specific circumstances (Martin & Hosken, 2003).

Furthermore, the evolution of Dobzhansky-Muller incompatibilities (DMIs) between populations is known to promote speciation. Neutral allelic substitution within a population can be incompatible with loci of a divergent population, and these incompatibilities are thought to be generated by various forms of genomic conflict (Seehausen *et al.*, 2014). Negative epistasis reduces the overall viability and sterility of their hybrids, acting as a powerful force underlying incipient reproductive isolation.

A powerful approach to understanding the strength and dynamics of postzygotic isolation is the study of hybrid zones, regions where divergent populations interbreed and produce offspring (Harrison, 1990; Harrison, 1993). A secondary hybrid zone emerges when two genetically and geographically distinct populations interbreed after expansion or migration (Jiggens & Mallet, 2000). One striking example of a secondary hybrid zone has been discovered in the Caribbean Islands and southeastern United States. In this region, two distinct populations of *D. melanogaster,* originating from west Africa and Europe (Kao *et al.,* 2014; Yukilevich et al., 2010), have recently come into secondary contact (Bergland *et al.,* 2014). After a migration event from Africa to current day Europe, these populations have been evolving in allopatry for approximately 10,000 to 15,000 years (Capy *et al.,* 1986). Secondary contact occurred in two waves, first with west African flies migrating to the Caribbean Islands during the transatlantic slave trade 400 to 500 years ago, and then the European flies arriving to the east coast US with European colonists < 200 years ago (Capy *et al.,* 1986; Duchen *et al.*, 2013).

Caribbean populations have peculiar morphological, behavioral, and pheromonal differences. They display exceptional African-like morphology based on body size, allozyme frequencies, hydrocarbon composition, and sequence variation (Kao et al., 2014; Capy *et al.,* 1986; Caracristi & Schlotterer, 2003; Yukilevich & True, 2008a). Sequence data suggest that United States flies display higher proportion of African alleles than do European flies, suggesting Caribbean populations as a potential source of African alleles introgression for North America populations (Kao et al. 2014; Yukilevich et al., 2010; Caracristi & Schlotterer, 2003; Yukilevich & True, 2008b; Capy *et al.,* 1986). Mating preferences and other premating/prezygotic reproductive barriers have been formally treated in this system showing partial sexual isolation between west African flies and American flies, but not Caribbean flies and male courtship behavior differing between American and Caribbean flies (Yukilevich & True 2008a, b). However, the presence of postmating sexual isolation in these American and Caribbean populations remains unexplored.

Our study aims to explore the role of postmating reproductive barriers in a *Drosophila melanogaster* secondary contact hybrid zone and to better understand how patterns of postmating barriers reflect the colonization history of fly populations in the area. We have investigated the role of remating, female egg laying, hatchability of laid eggs, and female longevity after mating with different males as putative postmating reproductive barriers. These phenotypes are good candidates for assaying the roles of extrinsic and intrinsic postmating reproductive barriers. We measured each of these phenotypes in females from different locations in the southeastern US and Caribbean islands to examine them for geographical patterns – which may reveal if and how these barriers affect this secondary contact zone of *Drosophila melanogaster* and by investigating these barriers, we provide insight into how these mechanisms of speciation function in a genetically admixed system.

## Materials and Methodology

### Fly Lines and Rearing Conditions

For our phenotypic assays, we used 23 isofemale lines of *Drosophila melanogaster* collected in the summer of 2004 and 2005 (Yukilevich & True, 2008b). The origins of the *Drosophila* are as follows (TABLE 1; FIGURE 1): Birmingham, AL (lines 1-1 and 1-2); Selba, AL (lines 2-1 and 2-2); Meridian, MS (lines 3-1 and 3-2); Thomasville, GA (lines 4-1 and 4-2); Tampa Bay, FL (lines 5-1 and 5-2); Sebastian, FL (line 6-1); Freeport, Grand Bahamas-west (lines 7-1 and 7-2); Bullock’s Harbor, Berry Islands (lines 8-1 and 8-2); Cockburn Town, San Salvador (lines 9-1 and 9-2); George Town, Exumas (lines 10-1 and 10-2); Mayaguana, Mayaguana (lines 11-1 and 11-2); Port Au Prince, Haiti (lines 12-1 and 12-2). Latitude and longitude coordinates can be found in Yukilevich and True (2008b). All flies were maintained at 25 °C in vials on a standard cornmeal diet (recipe available upon request) and entrained under a 12hr light:12hr dark regime.

**FIGURE 1.**
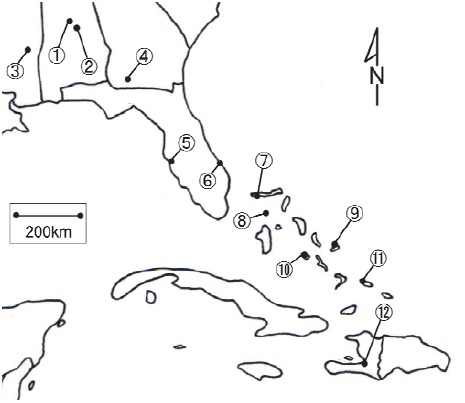
Map of locations used in postmating assays with numbers corresponding to those of Table 1.

**TABLE 1.** Locations, strain names, and line ID numbers of fly lines used in assays.

| Map Number | Location | Line(s) in order of decreasing latitude (N to S) | Line ID#'s (Yukilevich and True, 2008b) |
| --- | --- | --- | --- |
| 1 | Birmingham, AL | 1-1 and 1-2 | 21, 39 and 21, 36 |
| 2 | Selba, AL | 2-1 and 2-2 | 20, 28 and 20, 17 |
| 3 | Meridian, MS | 3-1 and 3-2 | 24, 2 and 24, 9 |
| 4 | Thomasville, GA | 4-1 and 4-2 | 13, 34 and 13, 29 |
| 5 | Tampa Bay, FL | 5-1 and 5-2 | 4, 12 and 4, 27 |
| 6 | Sebastian, FL | 6-1 | 28, 8 |
| 7 | Freeport, Grand Bahamas - West | 7-1 and 7-2 | 33, 16 and 33, 11 |
| 8 | Bullock's Harbor, Berry Islands | 8-1 and 8-2 | 40, 23 and 40, 10 |
| 9 | Cockburn Town, San Salvador | 9-1 and 9-2 | 42, 23 and 42, 20 |
| 10 | George Town, Exumas | 10-1 and 10-2 | 36, 9 and 36, 12 |
| 11 | Mayaguana, Mayaguana | 11-1 and 11-2 | 43, 19 and 43, 18 |
| 12 | Port Au Prince, Haiti | 12-1 and 12-2 | H, 29 and H, 25 |

### Egg laying, Hatchability, and Remating Rate Assays

Virgin females were collected from all 23 isofemale lines. Male flies up to one day old were collected from two lines (lines 1-2 and 11-1) located at polar ends of our geographical study region. We chose these two lines as sources for male flies based on clinal distance as well as maximal difference between courtship profiles and physical characteristics (Yukilevich & True, 2008b) to account for female mate preference, which has been previously established (Yukilevich & True, 2008a). All flies were collected on light CO2 anesthesia and aged for three to four days before entering our assays. We set up a full factorial experiment in which females from each of the isofemale lines were crossed with the two lines from which males were collected. Each cross was replicated 15 times.

All flies were live manipulated using aspirators for the remainder of the phenotypic assays to avoid any physiological and behavioral effects of CO_2_ anesthesia (Badre *et al.*, 2005). Assays lasted 24 days and were conducted in two stages. During the first 10 days (i.e. first stage) female remating rates and egg laying rates were measured over a 10-day period; during the following 14 days (i.e. second stage) hatchability rates were quantified.

In the first stage, females were transferred daily by aspirator into new vials with standard cornmeal fly food and blue food coloring. The dye helped visualize eggs laid by females without causing any variability in their behavior (Bergland, 2012). The vials also had 20 uL of a 10% diluted active yeast mixture to stimulate females’ reproductive activity. At lights on (i.e. dawn) on the initial day of the first stage, individual females were aspirated into a vial with two males from either one of the two male lines for mating. Approximately 90 minutes were allocated for copulation to occur, and all males were discarded immediately after this time period using an aspirator. Females that did not mate on the first day did not continue in the assay. Fecundity assays were conducted daily after the females were transferred into new vials. To assess short-term and long-term receptivity to remating effects, each individual female was introduced to two new males of the same genotype from her initial mating on the fourth and eighth day of the assay. Again we allowed 90 minutes on both remating days for copulations to occur, and all males were discarded via aspirator thereafter.

Incorrectly sexed vials in which the female - instead of the male - were accidentally discarded were not included in later analysis. Remaining vials that passed the first stage of the experiment were monitored daily for fly eclosion. Flies that eclosed were recorded and discarded immediately. Fly eclosion monitoring ended when either (a) three consecutive days of zero fly eclosions occurred or (b) 14 days of monitoring was reached - whichever occurred first. All phenotyping assays during the first and second stages were conducted within the first three hours of lights on (i.e dawn). All flies from the first stage and eclosing vials in the second stage were incubated at a controlled 25 °C with a light timer set for a 12hr light: 12hr dark regime.

### Longevity Assays

Female flies used in our longevity assays come from (arranged from north to south) Selba, Alabama, USA (line 2-2), Thomasville, Georgia, USA (line 3-1), Freeport, Grand Bahamas-west (line 7-2), Bullock’s Harbor, Berry Islands (line 8-1), and Port Au Prince, Haiti (line 12-2). Representative ‘American’ and ‘Caribbean’ males were derived lines originating from the same male collection lines used in egg laying, hatchability, and remating assays, Birmingham, Alabama, USA (line 1-2) and Mayaguana, Mayaguana (line 11-1), respectively. ‘Homotypic’ crosses were defined as male and female both of either American or Caribbean origin (i.e. American x American or Caribbean x American). “Heterotypic” crosses were defined as male and female from different origins (i.e. American x Caribbean or Caribbean x American). Males and females from the same origin were assumed to be more related and genetically similar to each other than those from different origins based on previous evidence (Yukilevich & True, 2008b).

Virgin females were collected on light CO2 anesthesia and aged singly in vials for four days. Males were collected in the same manner and aged in groups of five per vial. We performed crosses in two separate rounds, which lasted approximately 70 and 80 days. In the first round, we crossed female flies from Selba, Alabama, USA and Port Au Prince, Haiti to either our representative ‘American’ or ‘Caribbean’ male. There were 50 replicates for each unique cross. Because of the large effect size from our initial round, we had 25 replicates for each type of cross in the rest of our lines. In each round, aged female flies were placed with five male flies for 48 hours to ensure mating occurred. Male flies were discarded using an aspirator after the mating period. Female flies were then observed on a regular basis five days per week. Dates of deaths were recorded until the end of the 70 and 80-day observation periods. The females were transferred to fresh vials every seven days.

### Post-mating behavior data analysis

We examined the effects of geographic location on the total number of eggs laid by females, the total hatchability of those egg laid, and the propensity of females to remate three and seven days after initial mating day. For egg laying and hatchability, we used ANOVA to test the effects of latitude and longitudinal coordinates (i.e. geographic position effects). We also used the male and female identity and phenotyping blocks to account for the variation from genotypes of male and females in addition to experimental block effects.

Because remating was scored as a binary variable of whether or not the female copulated on the two remating days, we used logistic regression models to assess the effects of geographic location while controlling for male and female genotypes and block effects on short- and long-term female receptivity to remating. The significance of longitudinal and latitudinal coordinates and model fits were assessed using analysis of deviance tables.

We performed a permutation test to investigate the significance of the lower hatchability rates in the three central locations as revealed by logistic regression models as well as through visual confirmation of plots. We calculated the difference in hatchability between the five lines from our three central locations and the hatchability of all other fly lines (18 lines). We then randomly assigned fly lines into groups of 5 and 18 and calculated the difference in hatchability between these two groups. These permutations were repeated 10,000 times. P-values were calculated by the number of times the difference in hatchability between these two groups were equal to or greater than our observed value divided by our 10,000 permutations. The line with the lowest hatchability was removed for a follow-up permutation test to confirm that the lower hatchability was only due to the effect of one line. Similar permutation tests were conducted on total egg counts to determine that lower hatchability was also not due to lower egg counts via ascertainment bias. Hatchability of eggs laid by females mated to representative ‘American’ and ‘Caribbean’ males were performed separately, and P-values from these tests were corrected using the Bonferroni method.

All analyses were performed in R and the code for the permutation test is available upon request.

### Longevity data analysis

Survival analysis is used for temporal data of waiting times to an event with censored data. We employed methods from survival analysis to examine our data. We analyzed the waiting times of female death after homotypic or heterotypic mating. Females that escaped or survived past our observational periods were considered censored data points. The first step of survival analysis is to estimate survival functions for each of our crosses, S(t), which in our study is the probability of a female living longer than time, t. This can be done non-parametrically using the Kaplan-Meier method (Kleinbaum & Klein, 2012). Parametric models were tested (i.e. exponential, log-normal, log-logistic, and generalized gamma), but none yielded a good fit (data not shown). After survival curves were fitted, we used it to estimate the cumulative hazard function, H(t), for each type of cross. The cumulative hazard function shows the cumulative probability that a female has expired up to time, t.

The most common statistical test used for comparing survival distributions is the log-rank test. However, this test has the proportional hazards assumption, which requires that the hazard functions of the two groups being compared are parallel. Hazard functions for our comparisons of female longevity after heterotypic and homotypic matings were plotted and visually checked for the crossing of hazard curves. When hazard curves cross, the proportional hazards assumption is violated so another test must be conducted because the standard log-rank test has little to no power (Klein & Moeschberger, 1997). We chose to use a combined weighted log-rank test, which takes into account crossing hazard curves (Bathke *et al.*, 2005). This improved log-rank test has more power than the standard log-rank tests when the hazard functions cross and the hazard ratio is not proportional.

All analyses were performed in R using the ‘survival’ package to estimate the survival curves and hazard functions. The package ‘emplik’ was used as part of the improved log-rank test. The R code can be obtained online (http://www.ms.uky.edu/%7Emai/research/LogRank2006.pdf).

## Results

### Egg counts

Egg counts for each line were shown using side-by-side boxplots with locations arranged from the northernmost to the southernmost location, left to right (FIGURE 2A, 2B). It does not appear that egg counts follow a clinal or other geographical pattern for females mated to representative ‘American’ or ‘Caribbean’ males. There is much variation amongst the lines, but the median egg count for each location is approximately the same except for when the females from location 6 (line 6-1; Sebastian, FL) were mated to Caribbean males (FIGURE 2B). The ANOVA model showed that that most of the variance of egg laying was accounted for by male (p = 0.00167) and female (p<0.0001) genotypes as well as block effects (p < 0.0001) and that longitude and latitude were not significant influences (p = 0.32767, p = 0.49860). (SUPPLEMENTARY TABLE 1)

**FIGURE 2.**
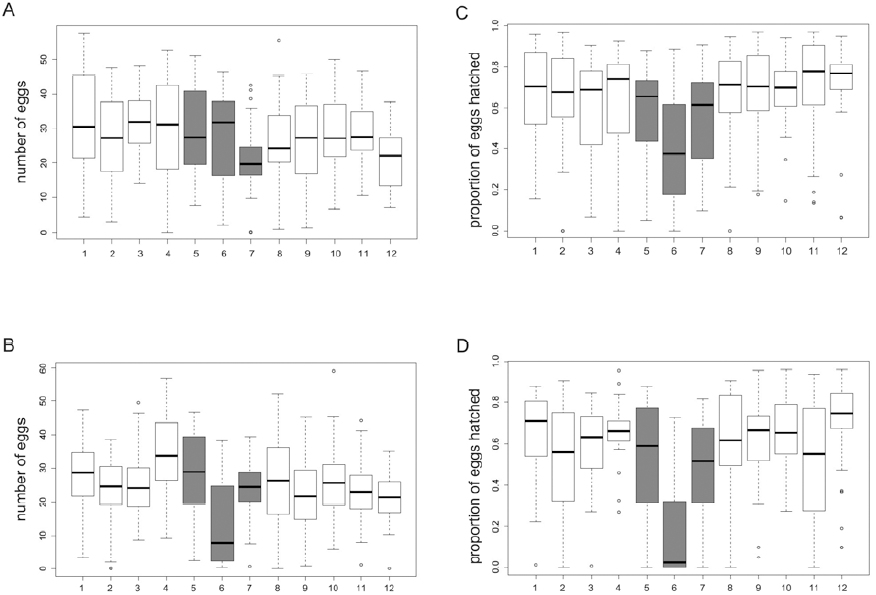
Egg counts of females mated with A) American males and B) Caribbean males. Hatchability of females mated with C) American males and D) Caribbean males. Each box plot is a isofemale line arranged from the northernmost location (left) to the southernmost location (right). Numbers on the X-axis correspond to those of Table 1.

### Remating

Short- and long-term remating rates for each isofemale line were plotted against latitude and longitude coordinates (SUPPLEMENTARY FIGURE 1, 2). Short-term remating rates were generally lower (range of rates: 0-30%) than long-term remating rates (range of rates: 0-60%). Remating rates do not appear to be influenced by location, which was investigated further with logistic regression.

The full logistic regression model evaluating effects of latitude and longitude while controlling for male and female genotypes and block effects found that latitude (p = 0.11) or longitude (p = 0.35) were not useful in predicting short-term remating rates with similar results for long-term remating rates (lon p = 0.7616, lat p = 0.6361). Male genotype was also not a significant influence on short-term or long-term remating rates (p = 0.4848 and p = 0.1240) (SUPPLEMENTAL TABLE 2, 4). The reduced models removing latitude and longitude as predictors showed that they were not significantly influencing remating rates (SUPPLEMENTAL TABLE 3, 5). Female identities in both logistic models for short- and long-term remating rates were significant, giving evidence that female genotypes could influence remating rates. However, when we fitted a model for long-term remating rates with a male x female interaction term, results showed that this interaction term was not significant (p = 0.0959) (SUPPLEMENTAL TABLE 6, 7).

### Hatchability

Hatchability for the various locations in the southeast US and Caribbean Islands were visualized using side-by-side boxplots with locations arranged from the northernmost to the southernmost location, left to right (FIGURE 2C, 2D). Hatchability in the three middle locations (location 4, 28, 33) at the border of the southeast US and Caribbean Islands appear lower than the locations on the edges in both the graphs displaying hatchability of females mated to American males (FIGURE 2C) and Caribbean males (FIGURE 2D).

Our ANOVA model took into account male and female identities and male/female genotype interactions on hatchability as well as experimental block effects while assessing influences of longitude and latitude (TABLE 2). Longitude had a significant effect on hatchability (F=3.954, p=0.472) while latitude did not (F=1.4, p = 0.2372) further suggesting that geographic location had some influence on hatchability rates as indicated by the dip in hatchability in FIGURE 2C, D.

**TABLE 2.** ANOVA table for hatchability model.

|  | DF | Sum Sq | Mean Sq | F value | P(>F) |
| --- | --- | --- | --- | --- | --- |
| Block | 14 | 2.934 | 0.2096 | 4.688 | <3.22e-08* |
| Female | 22 | 7.722 | 0.3510 | 7.853 | <2e-16* |
| Male | 1 | 0.887 | 0.8869 | 19.844 | 9.84e-06* |
| Latitude | 1 | 0.063 | 0.0626 | 1.400 | 0.2372 |
| Longitude | 1 | 0.177 | 0.1767 | 3.954 | 0.0472* |
| Female:Male | 22 | 1.298 | 0.0590 | 1.320 | 0.1493 |
| Residuals | 672 | 30.035 | 0.0447 |  |  |
| Residuals | 672 | 5097909 |  |  |  |

To evaluate the significance of the dip in hatchability rates, we performed permutation tests as described in our methods section. We found that the hatchability in the middle three locations was significantly lower than the rates in the surrounding locations regardless of the female being mated to an American male (p < 0.0001) or Caribbean male (p <0.0001). Results were similar when the location with the lowest hatchability rate was removed (28: Sebastian, FL, USA) and the permutation tests performed again (females mated to American male: p = 0.0056; females mated to Caribbean male: p = 0.0272). Similar tests were conducted on egg counts to investigate whether the lower hatchability was due to lower egg counts (i.e. ascertainment bias from not observing enough progeny). No significant differences in egg counts between females from the middle locations and the outer locations were found regardless of whether they were mated to American males (p = 0.3192) or Caribbean males (p = 0.7584). The same results were yielded when we removed the influence of the extremely low middle location, 28: Sebastian, FL, USA, (mated to American males: p=0.3016, mated to Caribbean males: p = 1.0). These results suggest a generalizable central location effect on hatchability.

### Longevity

Five female lines were measured for longevity after experiencing homotypic or heterotypic matings. The homotypic cross survival curves for females from lines 2-2, 3-1, and 12-2 were consistently higher than the survival curves of females in heterotypic crosses (FIGURE 3A, B, E). There were no apparent differences between homotypic and heterotypic survival curves of females originating from lines 7-2 or 8-1 (Figure 3C, D).

**FIGURE 3.**
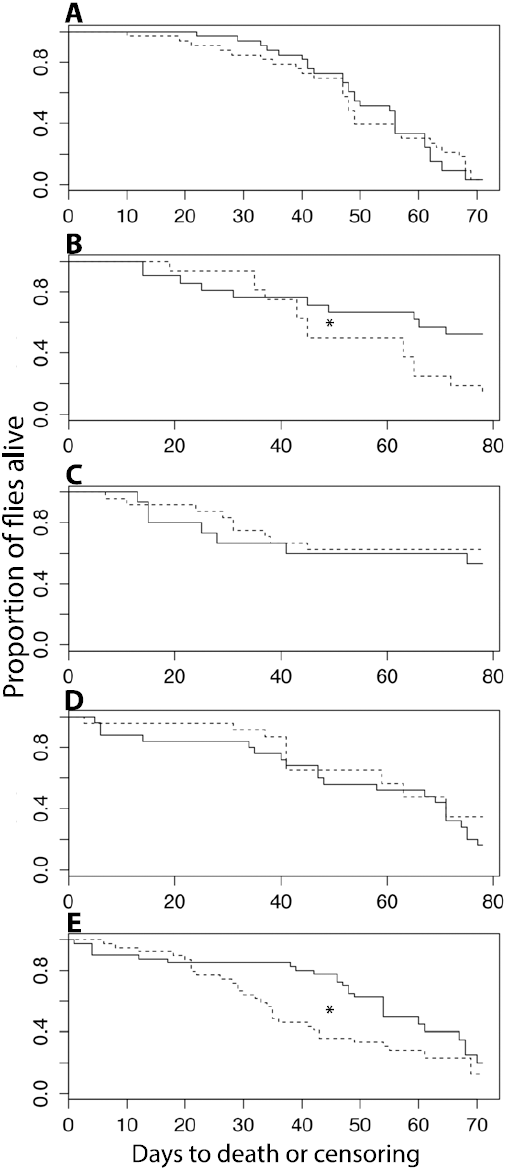
Survival curves of females of isofemale lines A) 2-2, B) 3-1, C) 7-2, D) 8-1, E) 12-2 after experiencing homotypic (solid line) or heterotypic (dashed line) matings. * indicates significant p-value < 0.05

**TABLE 3.**
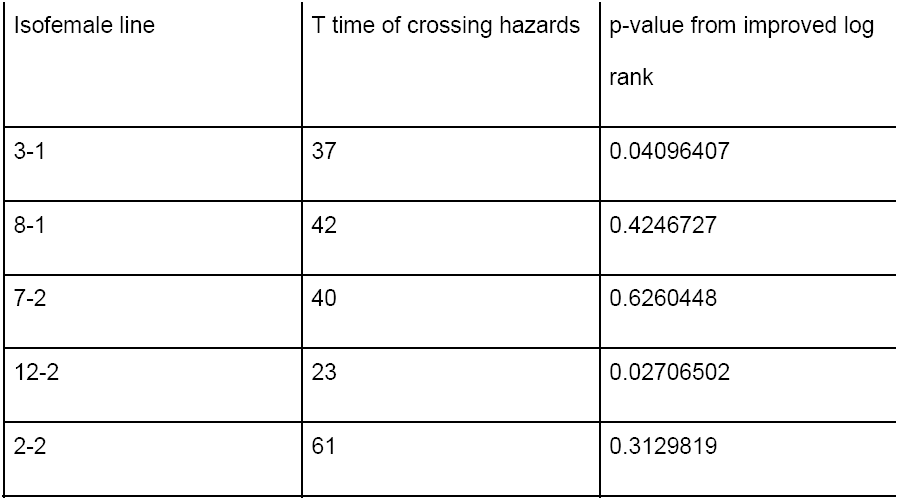
Improved Log-rank Test Results.

Hazard curves for all crosses and lines revealed non-proportional hazards in almost all cases of homotypic and heterotypic matings (SUPPLEMENTARY FIGURE 3). Crossing points of all hazard functions were visually estimated for use in the improved log-rank tests (TABLE 3). The improved log-rank tests showed evidence that females after heterotypic matings had shorter lifespans than females in homotypic matings for females from lines 3-1 and 12-2 (p = 0.0410 and p = 0.0271). Although, females of line 2-2 visually displayed a reduced lifespan when involved in heterotypic matings (FIGURE 3A), these results were not significant in our statistical test (p = 0.3130).

## Discussion

We examined several potential postmating reproductive barriers including remating rates, egg laying rates, hatchability, and female longevity that may potentially influence a system in the early stages of sexual isolation (Yukilevich and True 2008b). Our results illustrate the possible relationship between reproductive barriers and genetic admixture.

### Genetic admixture likely affects offspring fitness

We observed an interesting hatchability rate ‘valley’ produced by isofemale lines originating from our three central locations spanning the border of the United States and the Caribbean Islands (i.e. locations 5, 6, 7). These locations correspond to areas of high African and European admixture (Kao et al., 2014). This result may highlight the presence of essential genetic differences between American and Caribbean fly populations, which could have manifested as an intrinsic postzygotic barrier between these two populations. This type of evidence is indicative of the presence of Bateson-Dobzhansky-Muller incompatibilities (DMI), which are negative epistatic interactions and the most common form of intrinsic postzygotic isolation (Presgraves, 2010). A reduction in the fitness of ‘hybrid’ offspring here restricts the product of gene flow between American and Caribbean *D. melanogaster* populations. A more thorough investigation of these lines and genome sequences are required to confirm the presence of DMIs, but are beyond the scope of this study.

### Females evolve resistance to toxic males

We examined female longevity after mating with males that were more or less genetically related to them, as defined by physical distance, which does correlate with geographical distance (Kao et al., 2014). These results from the longevity assay were the inverse of our hatchability assays. Females originating from locations 7 and 8 did not seem as affected by heterotypic matings compared to females from the northern and southernmost locations (i.e. locations 2, 3, 12). It is known that male sperm has toxic effects on females after mating (Rice, 1996) and in response to this game of sexual conflict (at least in the laboratory), females develop ‘resistance’ against males that they coevolve with in the same environment (Arbuthnott *et al.*, 2014). Our findings not only support this coevolution tactic, but also illustrates that these patterns can naturally occur outside of the laboratory.

### Conclusions

Our study of postmating reproductive barriers along with previous investigations into premating barriers (Yukilevich & True 2008a, b) illustrate that pre- and post-mating barriers could be evolving at the same time and is not necessarily sequential. While our findings contribute to the ever-growing breadth of knowledge about sexual isolation and speciation, it also sheds light on the complexity of the interplay between isolating mechanisms and genetic admixture. Overall our data suggests that long-term postmating consequences - offspring fitness and female lifespan reduction - are of greater influence in this particular incipient sexual isolation scenario than when compared to more immediate postmating behavioral responses such as egg laying and remating receptivity. We have also observed the very possible effects of admixture at the border between the United States and Caribbean islands (i.e. locations 5, 6, 7) (Kao *et al.*, 2014) leading to interesting interactions between partially isolating mechanisms. Greater genetic admixture in flies originating from this area could promote the lower hatchability of eggs laid by females from these populations if American and Caribbean flies are genetically distinct enough to increase the possibility of DMIs occurring (Gompert *et al.,* 2012). The same genetic admixture could also contribute towards female hardiness against harm from mating with a wider range of genetically diverse males, which in turn can compensate for lower hatchability by increasing reproductive lifespan.

We did not find any evidence that egg laying rates or remating rates influenced the reproductive success in a systematic way with regard to these isofemale lines from the southeast United States and Caribbean Islands. However, the lack of evidence from our study does not imply that these behaviors in general are not influential postmating reproductive barriers. Current views of speciation regard the process as a sliding continuum in which speciation can move forward or step back and may even be arrested at intermediate stages (Seehausen *et al.*, 2014). Depending on the driving force of speciation, different types of reproductive barriers form at particular stages (Seehausen *et al.*, 2014). Thus, it may be that these postmating behaviors could be of importance at other stages in the speciation continuum, in which case, other species in the *Drosophila* genus may be better candidates to further investigate this question.

## Acknowledgements

We are grateful to R. Yukilevich for the isofemale lines from the southeast US and Caribbean Islands. We thank J. Saltz and B. Foley for useful discussions in designing behavioral assays. Additionally, we thank the numerous undergraduate research assistants including M. Hom, G. Kaur, M. Grozdanich, B. Kang, S. Walia, S. Ahmed, J. Ong, C. Kim, W. Liao, B. Mathews, L. Ishida, D. Kuo, and G. Liang who helped with the remating, egg laying, hatchability, and lifespan assays. We thank F. Zhang for piloting the lifespan assay as part of his high school summer internship. This work was supported by two NIH grants, GM102227 and MH091561. The authors of this paper have no conflicts of interests to declare.

